# miRBench: novel benchmark datasets for microRNA binding site prediction that mitigate against prevalent microRNA Frequency Class Bias

**DOI:** 10.1101/2024.11.14.623628

**Authors:** Stephanie Sammut, Katarina Gresova, Dimosthenis Tzimotoudis, Eva Marsalkova, David Cechak, Panagiotis Alexiou

## Abstract

**Motivation:** MicroRNAs (miRNAs) are crucial regulators of gene expression, but the precise mechanisms governing their binding to target sites remain unclear. A major contributing factor to this is the lack of unbiased experimental datasets for training accurate prediction models. While recent experimental advances have provided numerous miRNA-target interactions, these are solely positive interactions. Generating negative examples *in silico* is challenging and prone to introducing biases, such as the miRNA frequency class bias identified in this work. Biases within datasets can compromise model generalization, leading models to learn dataset-specific artifacts rather than true biological patterns.

**Results:** We introduce a novel methodology for negative sample generation that effectively mitigates the miRNA frequency class bias. Using this methodology, we curate several new, extensive datasets and benchmark several state-of-the-art methods on them. We find that a simple convolutional neural network model, retrained on some of these datasets, is able to outperform state-of-the-art methods. This highlights the potential for leveraging unbiased datasets to achieve improved performance in miRNA binding site prediction. To facilitate further research and lower the barrier to entry for machine learning researchers, we provide an easily accessible Python package, miRBench, for dataset retrieval, sequence encoding, and the execution of state-of-the-art models.

**Availability:** The miRBench Python Package is accessible at https://github.com/katarinagresova/miRBench/releases/tag/v1.0.0

## 1. Introduction

Over thirty years ago, the regulatory map of the animal cell was fundamentally changed when fragments of cellular RNA, previously considered non-functional, were identified as important regulators of the post-transcriptional life of RNA (Lee *et al*., 1993). MicroRNAs (miRNAs), small regulatory RNAs, were rapidly found to serve diverse roles, amongst which, functioning as master regulators of gene expression during embryogenesis, tissue development, and the maintenance of homeostasis in adults (Bernstein *et al*., 2003; Zhao *et al*., 2005; Ivey and Srivastava, 2010). Dysregulated miRNAs are linked to various diseases, including cancer (He and Hannon, 2004; Calin and Croce, 2006; Esquela-Kerscher and Slack, 2006), cardiovascular diseases (van Rooij *et al*., 2006; Ikeda *et al*., 2007; Thum and Condorelli, 2015), neurological disorders (Hébert and De Strooper, 2009), and immune conditions (O’Connell *et al*., 2007; Sonkoly and Pivarcsi, 2009; Dai and Ahmed, 2011). These molecules also hold promise as biomarkers (Condrat *et al*., 2020) and therapeutics (van Rooij and Olson, 2012; Rupaimoole and Slack, 2017; Chakraborty *et al*., 2017), including recent applications in CAR-T therapy (Rad *et al*., 2022; Shen *et al*., 2024). The importance of miRNAs in biology and biomedicine was highlighted with the Nobel Prize in Medicine in 2024 for their discovery.

miRNAs function by associating with proteins from the Argonaute (AGO) family, which are fundamental components of the RNA-induced silencing complex (RISC). In mammals, the four AGO proteins (AGO1-4) specifically interact with miRNAs to form ribonucleoprotein complexes central to RNA silencing. Among them, AGO2 is especially crucial; unlike AGO1, AGO3, and AGO4, its absence results in embryonic lethality in mice, highlighting its indispensable role (Liu *et al*., 2004; Morita *et al*., 2007). AGO2 identifies target RNA molecules by utilizing partial sequence complementarity between the ‘guide’ miRNA and the ‘target’ RNA (Bartel, 2004). In mammals most known AGO2 miRNA binding sites show partial complementarity, often focused on a ‘seed’ region located at the 5′ end of the ‘guide’ sequence. A ‘canonical seed’ sequence denotes a fully Watson–Crick complementary stretch of at least six nucleotides starting at the second position from the 5′ end of the miRNA ‘guide’. Further binding outside the seed area can stabilise the interaction and is commonly called 3’ compensatory binding. However, functional interactions not mediated by a ‘canonical seed’ have been known since the early days of miRNA targeting research in worms and mammals (Didiano and Hobert, 2006; Broughton *et al*., 2016). We, and others, have previously shown that less than 50% of experimentally identified miRNA binding sites are mediated by a canonical seed (Helwak *et al*., 2013; Klimentová *et al*., 2022; Hejret *et al*., 2023). While the exact rules of AGO2:miRNA target recognition remain largely unknown, computational prediction programs have been developed that rely on approximations of these rules, and form a crucial part of miRNA target gene prediction pipelines (Alexiou et al., 2009; Grešová et al., 2022). These programs commonly employ target site or ‘miRNA binding site’ prediction methods, which are not only vital components of larger pipelines but also essential tools for gaining deeper insights into the intricacies of AGO:miRNA:target interactions.

Several miRNA binding site classification models have been developed in the past few years with varying degrees of accuracy and generalizability (McGeary *et al*., 2019; Zheng *et al*., 2020; Min *et al*., 2022; Klimentová *et al*., 2022; Hejret *et al*., 2023; Yang *et al*., 2024). These models are based on various types of neural networks, including convolutional neural networks, residual networks, and attention networks as common architectures. Neural networks require substantial amounts of data to learn meaningful information related to the task at hand.

In 2013, an experimental method termed CLASH (Cross Linking, Ligation, and Sequencing of Hybrids) was developed (Helwak et al., 2013), utilising a ligation step between the ‘guide’ and ‘target’ sequences, produced ‘chimeric’ reads that contain both sequences of the AGO-bound guide:target interaction. More recently, another high-throughput technique termed chimeric eCLIP (enhanced Crosslinking and Immunoprecipitation) was developed (Manakov et al., 2022). The method integrates a chimeric ligation step into the eCLIP method, which has an improved library generation efficiency relative to earlier CLIP methods (Van Nostrand et al., 2016). Chimeric eCLIP is therefore also able to capture AGO-bound chimeric reads of guide and target RNA sequences, but at an even higher resolution and sensitivity, providing a comprehensive map of miRNA binding sites. An important distinction between the two methods is that in the CLASH method, AGO protein is induced, whereas chimeric eCLIP leverages endogenous AGO protein. Consequently, CLASH may be retrieving more interactions of lower affinity due to the higher abundance of AGO in CLASH compared to chimeric eCLIP, as we have previously suggested (Hejret et al., 2023). The majority of miRNA binding site classification models are trained on datasets derived from experiments that produce chimeric miRNA-binding site interactions (Helwak et al., 2013; Klimentová et al., 2022; Hejret et al., 2023), or on databases of experimentally derived binding sites (Vlachos et al., 2015; Chou et al., 2016; Pla et al., 2018). One model (McGeary et al., 2019) is instead trained on RNA Bind-n-Seq data, which yields a continuous estimation of dissociation constants that can serve as a proxy for binding affinity. Importantly, while these datasets predominantly contain experimentally validated target sites, representing “positive interactions,” machine learning approaches require a comparable number of “negative examples” for effective training. However, experimentally confirming miRNA *non*-binding sequence pairs is significantly more challenging. Consequently, negative examples are often underrepresented in database-derived datasets and are typically generated in silico for chimeric miRNA-binding site interaction datasets.

In this study, we uncover what we term a *miRNA frequency class bias* in existing miRNA binding site datasets used for training and benchmarking (Pla *et al*., 2018; Hejret *et al*., 2023). This bias occurs when the frequency distribution of miRNAs in the negative class differs from that in the positive class, arising from how negative examples are generated *in silico*. The use of such datasets for training on the task of miRNA binding site prediction results in models that struggle to generalize well, as they learn sequence interaction patterns muffled by the intricacies of the data.

To mitigate this bias, we developed a novel strategy for generating negative examples. We applied this strategy to existing datasets to create new, unbiased versions, and also curated a new dataset based on human AGO2 chimeric miRNA binding site interactions from a recent experimental method (Manakov *et al*., 2022). This new dataset, processed using our negative example generation strategy, yielded a training set containing over 2.5 million ∼1:1 class-balanced miRNA-binding site interactions. Alongside this large training set, we also generated smaller benchmarking datasets, all carefully constructed to mitigate the miRNA frequency class bias. We demonstrate that models trained on these unbiased datasets, particularly the large Manakov2022-derived training set, generalize better and outperform the current state-of-the-art on these benchmarks.

We make these new training and benchmarking datasets publicly available and provide a user-friendly Python package for easy access to these datasets, as well as to the state-of-the-art models benchmarked in this work.

## 2. Methods

### 2.1 Identification of *miRNA frequency class bias* in published datasets

To investigate potential imbalances in miRNA frequency between positive and negative classes in widely used miRNA binding site interaction datasets, we performed a series of analyses using simple classification models. We hypothesized that such imbalances, which we term “miRNA frequency class bias,” could lead to artificially inflated performance metrics for models trained on these datasets.

Two independent, publicly available datasets were used:

a. **Original_Hejret2023:** This dataset was obtained from the HybriDetector repository (accessed on 16-Dec-2024). It includes a class-imbalanced (1:10) training set^1^ and a class-balanced (1:1) test set^2^.
b. **miRAW:** This dataset was retrieved from the repository associated with (Pla et al., 2018)^3^, using the class-balanced (1:1) training set^4^ and test set^5^.

For each dataset, we extracted the mature miRNA sequences from each example. We then computed k-mer counts (k=3) from these sequences, generating a feature vector for each miRNA. These k-mer counts served as the sole input features for our classification models, allowing us to isolate the effect of miRNA frequencies. We trained a decision tree classifier on the training set of each dataset, using the miRNA k-mer counts as features and the original class labels (positive or negative) as targets. The trained models were then evaluated on their corresponding test sets. As a baseline, we also implemented a random classifier that assigns uniformly distributed values between 0 and 1 to each example, representing random guessing.

We evaluated model performance using the Average Precision Score (APS) instead of the Area Under the Precision-Recall Curve (AUPRC). While AUPRC is a common metric, it can be susceptible to overestimating performance when dealing with skewed score distributions or models that produce a limited range of prediction scores. APS, in contrast, provides a more robust measure of performance in such cases. It is calculated as the weighted mean of precision values at each prediction threshold, where the weights are the differences in recall between consecutive thresholds. Formally, APS is given by:

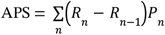

where *P*_*n*_ and *R*_*n*_ are the precision and recall at the nth threshold.

We observed in our preliminary analyses, particularly when evaluating the simple decision tree classifiers, that the distribution of prediction scores could be highly skewed. In such scenarios, a single high-scoring prediction followed by a large number of predictions with similar but slightly lower scores can artificially inflate the AUPRC. This is because the AUPRC calculation involves linear interpolation between points on the precision-recall curve. A large jump in recall, caused by many predictions having similar scores, can lead to an overestimation of the area under the curve. To avoid this potential overestimation and ensure a fair comparison across all models, we opted to use APS as the primary performance metric throughout this study.

### 2.2 Curating datasets corrected for *miRNA frequency class bias*

Publicly available data from high depth profiling of miRNA targets with the new chimeric eCLIP experimental method described by (Manakov *et al*., 2022), was downloaded from the Gene Expression Omnibus under SubSeries GEO ID GSE198250 (on 14-May-2024). The samples selected for download were to include only chimeric eCLIP data from the human cell line HEK293T. Only files for the R1 read sequences were used, yielding one file per sample to be processed. The files were selected using the GEOparse Python package (version 2.0.4) and downloaded using enaBrowserTools available on GitHub^6^.

The downloaded samples were pre-processed as per part of the pipeline for the analysis of total chimeric eCLIP datasets described by (Manakov *et al*., 2022). The utilized part of the pipeline, available on their GitHub, trims the 3’ adapters, and trims 10 nucleotides from the 3’ end to ensure no random sequences from the 3’ UMI remain.

The sample files were then processed by HybriDetector (Hejret *et al*., 2023), a chimeric read annotation pipeline available on GitHub^7^, to filter and separate different types of reads. The pipeline outputs high confidence pairs of guide (non-coding RNA) to target (binding site) sequences. To handle larger files from (Manakov *et al*., 2022), some modifications were made to the original pipeline, and have been made available on GitHub^8^. A total of 19 sample files were successfully processed with HybriDetector. The files were vertically concatenated into a single dataset containing all the examples of the positive class and made publicly available on Zenodo^9^.

Two corresponding, publicly available datasets, from (Klimentová *et al*., 2022; Hejret *et al*., 2023), also processed using the HybriDetector pipeline, and each containing the positive class curated from data derived from chimeric eCLIP and CLASH experimental methods, respectively, were downloaded from GitHub^10^ and ^11^, respectively, producing a total of three positive class datasets (Klimentova2022, Hejret2023, and Manakov2022), for each of which the negative class was generated *in silico*.

These three positive class datasets were processed via a series of post-processing pipelines available on GitHub^12^, primarily to generate the negative class and create the train/test splits.

The datasets were first filtered to retain only sequence pairs for which the non-coding RNA is a miRNA, eliminating other annotated non-coding RNA types, such as tRNAs and yRNAs. Next, the datasets were deduplicated based on the combination of miRNA to binding site sequence, to ensure unique positive examples. Positive examples with miRNA families unique to the Manakov2022 dataset were extracted and labelled as the Manakov2022 left-out dataset. This separate dataset is useful for evaluating the generalisability of models trained on any of the train sets presented here.

Next, a series of steps were performed to generate the negative examples. For each of the datasets, unique binding site sequences with >90% similarity were clustered together. Then, to prevent the bias identified in previous datasets, negatives were generated per miRNA family in a way that retains the same proportion of miRNA families in each of the positive and negative classes. The dataset was alphabetically sorted by the miRNA family name.

For each group of positive examples with miRNAs in the same miRNA family, negative examples with binding sites from clusters other than the ones in those positive examples were selected. These candidates were randomly sampled, allowing only one binding site per cluster to be included in the negative examples for that miRNA family group. The described procedure ensures that miRNA binding site pairs in the negative class are sufficiently different from those in the positive class. A positive to negative class ratio of 1:1 was produced for each dataset.

The Hejret and Manakov datasets were then split into train and test sets according to which chromosome the binding site sequence maps to. Binding site sequences from chromosome 1 were assigned to the test set and the rest were assigned to the train set, to ensure binding sites in the train and test sets were sufficiently dissimilar.

This series of post-processing pipelines generated a total of 6 comprehensive datasets; Klimentova2022 test set (477 positives), Hejret2023 train (4,084 positives) and test (495 positives) sets, and Manakov2022 train (1,253,320 positives), test (168,342 positives), and left-out (10,027 positives) sets consisting of miRNAs not found in any of the other datasets.

### 2.3 State of the Art: miRNA binding site prediction models

**TargetScanCnn_McGeary2019** (McGeary *et al*., 2019) is a convolutional neural network with two convolutional layers and two fully connected layers. It uses an outer product of a one-hot-encoded representation of the first 10 nucleotides of a miRNA and 12 nucleotides of its putative target. All possible 12-nucleotide subsequences of the putative target are scored and the highest score is used as a final prediction. TargetScanCnn was trained using RNA Bind-n-Seq (RBNS) –derived dissociation constants (K_d_) and mRNA-transfection fold-change measurements produced by the authors. The model predicts –log(K_d_) between miRNA and putative target, which can be used as a proxy for a binding affinity score.

**CnnMirTarget_Zheng2022** (Zheng *et al*., 2020) uses a convolutional neural network consisting of four convolutional layers with max-pooling followed by two dense layers with dropout. It uses a one-hot-encoded representation of miRNA concatenated with a putative target, padded to 110 nucleotides. CnnMirTarget was trained on data from three sources: 1) human AGO1 CLASH Helwak 2013 data (Helwak *et al*., 2013); 2) *C. elegans* and mammalian iPAR-CLIP data (Grosswendt *et al*., 2014); and 3) MirTarBase data with strong experimental evidence (Hsu *et al*., 2011). The model’s predictions may be directly used to measure the probability of binding.

**TargetNet_Min2021** (Min *et al*., 2022) is a deep learning model based on the ResNet architecture (He *et al*., 2015). It uses a one-hot-encoded representation of a miRNA and its putative target after extended seed alignment. TargetNet was trained on the miRAW dataset (Pla *et al*., 2018). This dataset was constructed from four sources: 1) Diana TarBase (Vlachos *et al*., 2015); 2) MirTarBase (Chou *et al*., 2016); 3) human AGO1 CLASH Helwak 2013 data (Helwak *et al*., 2013); and 4) *C. elegans* iPAR-CLIP data (Grosswendt *et al*., 2014). The model’s predictions may be directly used to measure the probability of binding. **miRBind_Klimentova2022** (Klimentová *et al*., 2022) is a deep learning model based on the ResNet architecture (He *et al*., 2015). It uses a two-dimensional representation of miRNA and putative target sequence. To address the imbalance between positive and negative classes, miRBind uses an instance hardness-based label smoothing approach, which keeps important, discriminative examples in the final training dataset and discards easily classifiable examples to rebalance the skewed ratio (Guo and Viktor, 2004). miRBind was trained on AGO1 CLASH Helwak 2013 data (Helwak *et al*., 2013). The model’s predictions are directly used to measure the probability of binding. **miRNA_CNN_Hejret2023** (Hejret *et al*., 2023) is a convolutional neural network consisting of six convolutional layers followed by dense layers. It uses a two-dimensional representation of miRNA and putative binding site interaction introduced in miRBind (Klimentová *et al*., 2022). miRNA_CNN_Hejret2023 was trained on the AGO2 CLASH Hejret 2023 training set that is also presented in this manuscript. The model’s predictions are directly used to measure the probability of binding.

**InteractionAwareModel_Yang2024** (Yang *et al*., 2024) is a deep multi-head attention network consisting of three parts: 1) sequence feature extraction; 2) interaction pattern identification; and 3) classification. It uses a one-hot-encoded representation of miRNA sequence padded to 30 nucleotides and the first 40 nucleotides of a putative target. The InteractionAwareModel was trained on re-analyzed data from human AGO1 CLASH Helwak 2013 (Helwak *et al*., 2013). This model’s predictions may also be used directly to measure the probability of binding. Beyond these ML methods that have been specifically trained for the task, we also implemented two simple methods based on co-folding and seed.

**RNACofold** is a tool offered by the ViennaRNA Package (Lorenz *et al*., 2011). It computes the binding energy and base-pairing pattern of an input pair of interacting RNA molecule sequences. The model’s output predicted minimum free energy of the folded structure is multiplied by –1 to be used as a measure of binding affinity.

For the seed measure, we followed previous studies (McGeary *et al*., 2022) and four different definitions of seed were used: 1) **Seed8mer** –full complementarity on positions 2-8 and A on position 1; 2) **Seed7mer** –full complementarity on positions 2-8, or full complementarity on positions 2-7 and A on the position 1; 3) **Seed6mer** –full complementarity on positions 2-7, or full complementarity on positions 3-8 or full complementarity on positions 2-6 and A on the position 1; and 4) **SeedNonCanonical** –the same as Seed6mer, but allowing for a single bulge or mismatch.

### 2.4 Retraining the miRNA_CNN_Hejret2023 model

We retrained the miRNA_CNN_Hejret2023 model on some of the new bias-corrected datasets presented here. To evaluate the effect of correcting the miRNA frequency class bias on the performance of the miRNA_CNN_Hejret2023 model, we retrained the model on the corrected Hejret2023 train set, to compare its performance with that of the original model that was trained on the (biased) Original_Hejret2023 dataset.

Next, we investigated the reliance of the performance of the model on dataset size. We retrained the miRNA_CNN_Hejret2023 model on 5 datasets of increasing sizes, subsampled from the Manakov2022 train set, and on the full Manakov2022 train set. The subsampling was done randomly while keeping the positive:negative ratio exactly 1:1. The number of positives in the training datasets are: 100, 1,000, 4,084 (equivalent to the number of positives in the bias-corrected Hejret2023 train set presented here), 10,000, 100,000, and 1,253,320 (the full Manakov2022 train set). Retraining of the miRNA_CNN_Hejret2023 model for all dataset sizes was carried out in the same way as in the original paper including all the hyperparameter settings (Hejret *et al*., 2023).

## 3. Results and Discussion

### 3.1 microRNA frequency class bias can affect target site classification performance

A critical issue hindering accurate miRNA binding site prediction is the *microRNA frequency class bias*. This bias arises when certain miRNAs appear disproportionately more often in the positive class than in the negative class of published datasets. This skewed representation is often exacerbated by the common practice of generating negative examples by pairing random miRNAs with binding sites. Consequently, highly abundant miRNAs in the positive class are often underrepresented in the negative set..

To investigate the impact of this issue, we conducted a sanity test using a previously published (Hejret et al., 2023) chimeric read dataset (Original_Hejret2023). We used the train set to train a simple decision tree machine learning model using only the miRNA sequence as input, excluding any binding site information and effectively isolating the influence of miRNA frequencies. This model cannot learn anything about miRNA binding rules as it is missing information about the miRNA binding site interaction. We would therefore expect this model to perform no better than random chance (Average Precision Score (APS) ∼ 0.5). However, the reported APS on the Original_Hejret2023 test set was much higher (APS ∼ 0.75). A similar analysis on another widely used dataset (Pla et al., 2018), which also exhibits this miRNA frequency class bias, produced the same effect, with the decision tree model trained only on miRNA sequences significantly outperforming random chance (APS ∼ 0.85).

These findings confirm that even simple models can exploit miRNA frequency imbalances between classes in the data, achieving deceivingly high predictive performance on the training data, but failing to generalize to new datasets. This miRNA frequency imbalance allows the models to learn spurious patterns in the miRNA frequency distribution rather than the true underlying features of the data such as sequence complementarity, structural accessibility, or legitimate miRNA binding rules. This artifact poses a significant threat to model generalization. If a classifier relies on the disproportionate presence of certain miRNAs in the positive class, rather than actual binding patterns in the miRNA binding site interaction, it may fail to accurately predict miRNA binding sites when tested on a dataset featuring a different miRNA frequency distribution.

### 3.2 Alternative negative example generation method can correct the miRNA frequency class bias

We have proceeded to correct the microRNA frequency class bias in one of the previously published datasets (Hejret et al., 2023) by implementing a novel procedure for negative example generation (Section 2.2). Briefly, instead of associating a random miRNA to each positive binding site, we attach a carefully selected binding site to each instance of a miRNA that occurs in the positive class, effectively ensuring a perfect match in miRNA frequency between the positive and negative classes across the entire dataset. When we attempted to train another simple decision tree classifier, using only the miRNA sequence as input, on the new corrected Hejret2023 dataset, the performance (APS=0.39) was lower than that of a random model (APS=0.50). This demonstrates that our method of negative example generation successfully addresses the previously identified artifact. The slight underperformance, however, requires further explanation. It arises because, while the entire dataset maintains a perfect balance of miRNA frequencies between positive and negative examples, the train/test split strategy does not explicitly enforce this balance. Consequently, random chance can lead to slight imbalances in miRNA frequencies between the training and testing sets. For example, the training set might, by chance, have slightly more positive examples for a particular miRNA than negative examples. Since our simple decision tree can only learn from miRNA sequences and has no access to binding site information, it might learn this incidental imbalance in the training set and predict “positive” more often for that miRNA. However, because the overall dataset is balanced, this slight overrepresentation in the training set will necessarily be mirrored by an underrepresentation in the test set. Therefore, the model’s learned bias from the training set will lead it to perform *worse* than random on the test set, explaining the observed APS of 0.39. It is important to note that this underperformance is a consequence of the limitations of the simple decision tree model and the train/test split strategy and does not invalidate the effectiveness of our negative example generation method in correcting the miRNA frequency class bias.

Having addressed the miRNA frequency bias, we considered the possibility of inadvertently introducing a similar bias related to the *binding site sequences* themselves. To investigate this, we trained another simple decision tree classifier, this time using only the binding site sequences as input and excluding any miRNA sequence information. This model achieved an APS of 0.50, comparable to the performance of a random classifier. This result suggests that our negative example generation procedure does not introduce a notable *binding site* frequency bias between the positive and negative classes. Therefore, we consider this procedure suitable for generating the negative class in miRNA:binding site datasets, promoting more robust training and benchmarking of models.

### 3.3 Machine learning prediction models are affected by microRNA frequency class bias

We proceeded to examine whether previously published miRNA binding site prediction models were affected by this artifact. We compared the APS of seven miRNA binding site prediction tools on the uncorrected Original_Hejret2023 dataset and the bias-corrected version (Hejret2023) presented here. This experiment demonstrates that several models are affected by the artifact, showing a notable drop in performance when evaluated in the corrected dataset (Table 1).

**Table 1.** Average Precision Score (APS) for several miRNA binding site prediction tools evaluated on the Original_Hejret2023 test set, and on the bias-corrected Hejret2023 test set presented here.

| Tool | (Original)<br>Hejret2023 test | (Corrected)<br>Hejret2023 test |
| --- | --- | --- |
| TargetScanCNN | 0.74 | 0.71 |
| CnnMirTarget | 0.63 | 0.53 |
| TargetNet | 0.66 | 0.58 |
| miRBind | 0.87 | 0.80 |
| miRNA_CNN_Hejret | 0.89 | 0.77 |
| Interaction Aware<br>Model | 0.77 | 0.74 |
| RNACofold | 0.77 | 0.74 |
| Random | 0.49 | 0.51 |

As expected, the model that was trained on the Original_Hejret2023 dataset, miRNA_CNN_Hejret2023, shows a large drop in performance when the artifact is corrected (APS: 0.89→0.77). Our ‘negative control’ model, RNACofold, which is not trained on any miRNA binding site interaction datasets, and therefore should not be affected by microRNA frequency class bias, shows a random variation drop in APS of 0.03 (APS: 0.77→0.74) similar to the InteractionAwareModel (APS: 0.77→0.74) and the TargetScanCNN (APS: 0.74→0.71), while a completely random model shows a random increase of similar magnitude (APS: 0.49→0.51). Surprisingly, we also notice that some other models show a drop in performance (miRBind APS: 0.87→0.80; TargetNet APS: 0.66→0.58; CnnMirTarget APS: 0.63→0.53). This is surprising because these models were trained and published before the Original_Hejret2023 dataset was produced in 2023, and thus could not have been directly affected by the microRNA frequency class bias we have detected in this dataset. However, we notice that all of these models have been fully or partially trained on an older AGO1 CLASH dataset (Helwak et al., 2013), which originates from an experiment that also happens to be performed on the same cell line (HEK293) as the Hejret dataset, and thus contain similar microRNA frequency class bias as the newer Original_Hejret2023 dataset. Therefore, the fact that models trained on other datasets also experienced a notable drop in performance, strongly indicates that the miRNA frequency class bias is a pervasive problem in the datasets currently available for miRNA binding site prediction model training. The widespread nature of this bias underscores the critical need for the development of corrected datasets, such as those presented in this study, to enable the training of accurate and generalizable miRNA binding site prediction models.

### 3.4 Novel miRNA binding site datasets

#### 3.4.1 Curation of datasets

Having shown that the majority of current state-of-the-art methods are affected by artifacts produced from a family of related training/testing datasets, and recognising the potential for other such datasets to introduce similar artifacts to future methods, we have produced a set of novel datasets with corrected microRNA frequency class bias. We curated three miRNA binding site datasets; Hejret2023, Klimentova2022, and Manakov2022, produced by two different high-throughput experimental protocols (CLASH, and chimeric eCLIP), and standardized through a post-processing pipeline (Section 2.2). The Hejret2023 and Klimentova2022 datasets are based on previously published datasets used in the training and/or evaluation of miRNA binding site prediction models (Hejret *et al*., 2023; Klimentová *et al*., 2022). The Manakov2022 dataset is completely novel, orders of magnitude larger than the others combined, and, to our knowledge, has not yet been used by any model to date for training or evaluation. To further ensure that future models using these datasets generalise better, we initially extract a ‘left-out’ set from the Manakov2022 dataset. The ‘left-out’ set contains only miRNAs from miRNA families that are unique to the Manakov2022 dataset, and not found in any of the Hejret2023 and/or Klimentova2022 datasets. The ‘left-out’ dataset is a small dataset, but it is crucial to ensure the generalizability of models trained on the Hejret2023 or Manakov2022 train sets. The remaining Manakov2022 dataset contains miRNAs from miRNA families that are also found in the Hejret2023 and/or Klimentova2022 datasets. We applied the negative example generation procedure (Section 2.2), that preserves the relative abundance of miRNAs in the positive and negative classes, to each of the Manakov2022 ‘left-out’ set and the remaining Manakov2022 dataset, and further split the latter into the ‘train’ and ‘test’ sets presented here.

#### 3.4.2 Programmatic access to datasets, encoders, and predictors

Adhering to FAIR principles, and in order to facilitate the availability and interoperability of the novel datasets presented here, we have developed a Python package called miRBench that allows easy access to all produced datasets. The latest version of the package is freely available on GitHub and has a permanent version deposited on Zenodo. The Python package is also distributed through the Python Package Index (PyPI) so that it can be installed using a single-line command, and can provide access to the datasets using only a single line of Python code. All data and code is publicly available (See Data availability).

The package is structured into three modules: dataset, encoder, and predictor. The dataset module is responsible for access to the benchmark datasets described in this paper. The main function of the module is `get_dataset_path(dataset_name, split, force_download)` which downloads the selected dataset of a specified split, from Zenodo, into the local cache, and returns the local path to the dataset. A complementary function is `get_dataset_df(dataset_name, split, force_download)` which instead of returning the path to the dataset, returns the dataset loaded into a Pandas DataFrame.

The encoder module is responsible for encoding data into the format expected by a predictor module. The main function of the module is `get_encoder(predictor_name)` which returns an instance of an encoder object implemented for a specified predictor. The encoder expects data as a Pandas DataFrame with columns named ‘noncodingRNA’ and ‘gene’. Specifying custom column names is possible when calling the encoder. The returned data format differs for every encoder and is specific to the predictor.

The predictor module is responsible for predicting miRNA-binding site interaction. The main function of the module is ‘get_predictor(predictor_name)‘ which returns an instance of a specified predictor object. The predictor object expects data encoded by a corresponding encoder and returns an array of predictions. Supported encoders and predictors in miRBench version 1.0.0 include TargetScanCnn_McGeary2019, CnnMirTarget_Zheng2020, TargetNet_Min2021, miRBind_Klimentova2022, miRNA_CNN_Hejret2023, InteractionAwareModel_Yang2024, RNACofold, Seed8mer, Seed7mer, Seed6mer, and Seed6merBulgeOrMismatch, described further in Section 2.3.

#### 3.4.3 Descriptive characteristics of positive samples in novel datasets

As part of the curation process of the three collections of datasets described, the target binding site sequences were aligned to the human genome via HybriDetector (Hejret et al., 2023), enabling assignment of genic feature annotations (Fig. 2A). The Klimentova2022 shows only 23% of binding sites mapping to regions annotated as 3’UTR. The other two datasets (Hejret2023 and Manakov2022) show a consistent prevalence of 3’UTR binding sites at 37% and 41% of annotated interactions, respectively. We observe that, overall, chimeric eCLIP appears to have more intronic targets than CLASH, at least for our limited dataset size.

**Figure 1:**
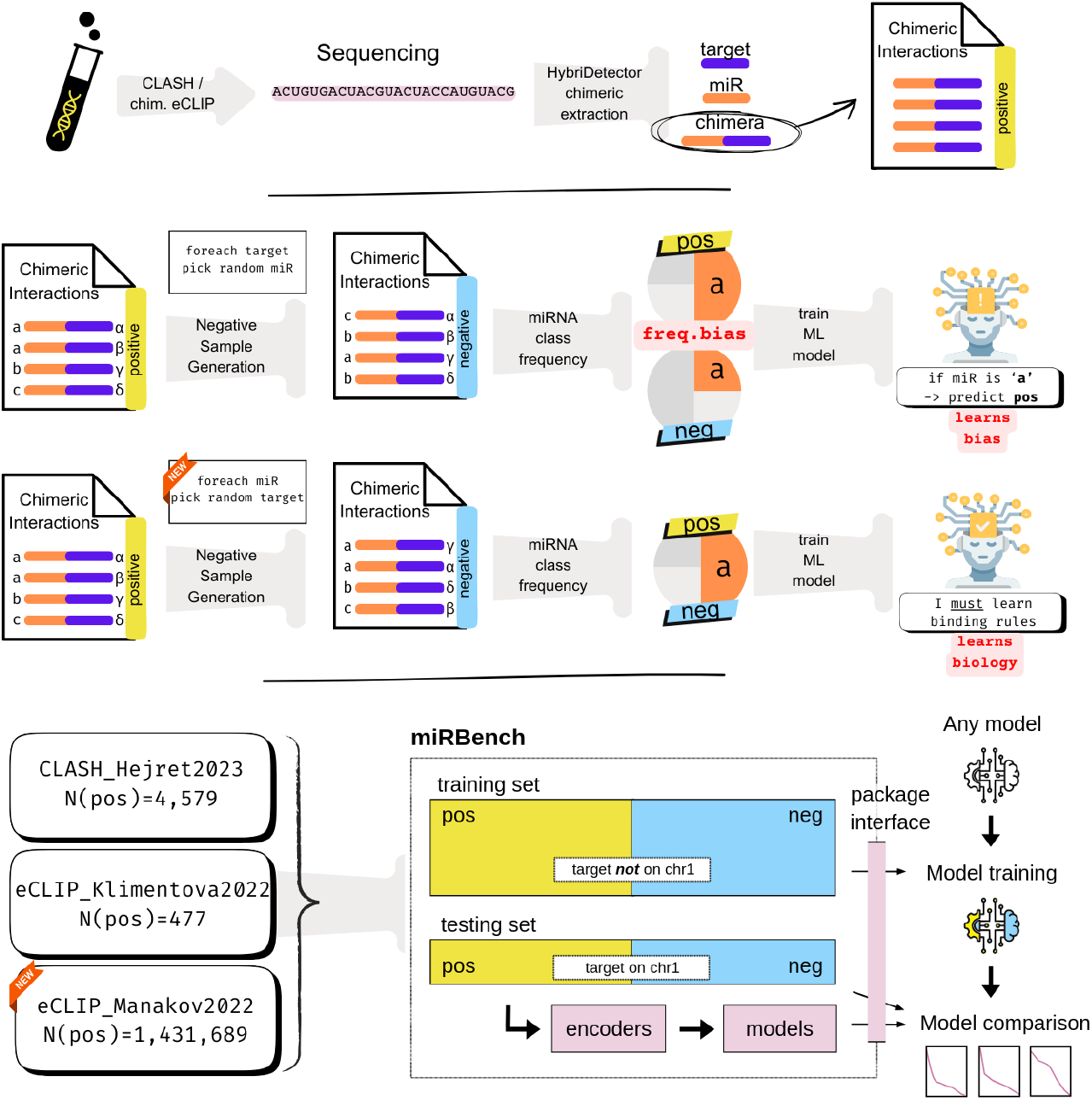
Graphical abstract. High-throughput experiments like CLASH and chimeric eCLIP identify miRNA-target interactions by sequencing chimeric reads containing fragments of both miRNA and target mRNA sequences. These chimeras form the basis for “positive example” datasets used in training miRNA target prediction models. However, generating corresponding “negative example” datasets is crucial. The conventional approach of pairing target sequences with random miRNAs (top) introduces a miRNA frequency class bias, leading machine learning models to learn spurious dataset-specific patterns rather than true miRNA-target binding rules. Our novel approach (bottom) pairs each miRNA with a selected non-cognate target sequence, removing this bias and forcing models to learn genuine binding rules. The miRBench Python package provides access to three curated datasets generated using this unbiased methodology, pre-split into training and testing sets. It also includes encoders for data formatting and implementations of state-of-the-art prediction models, facilitating robust training and benchmarking for the development of more accurate miRNA target prediction tools

**Figure 2.**
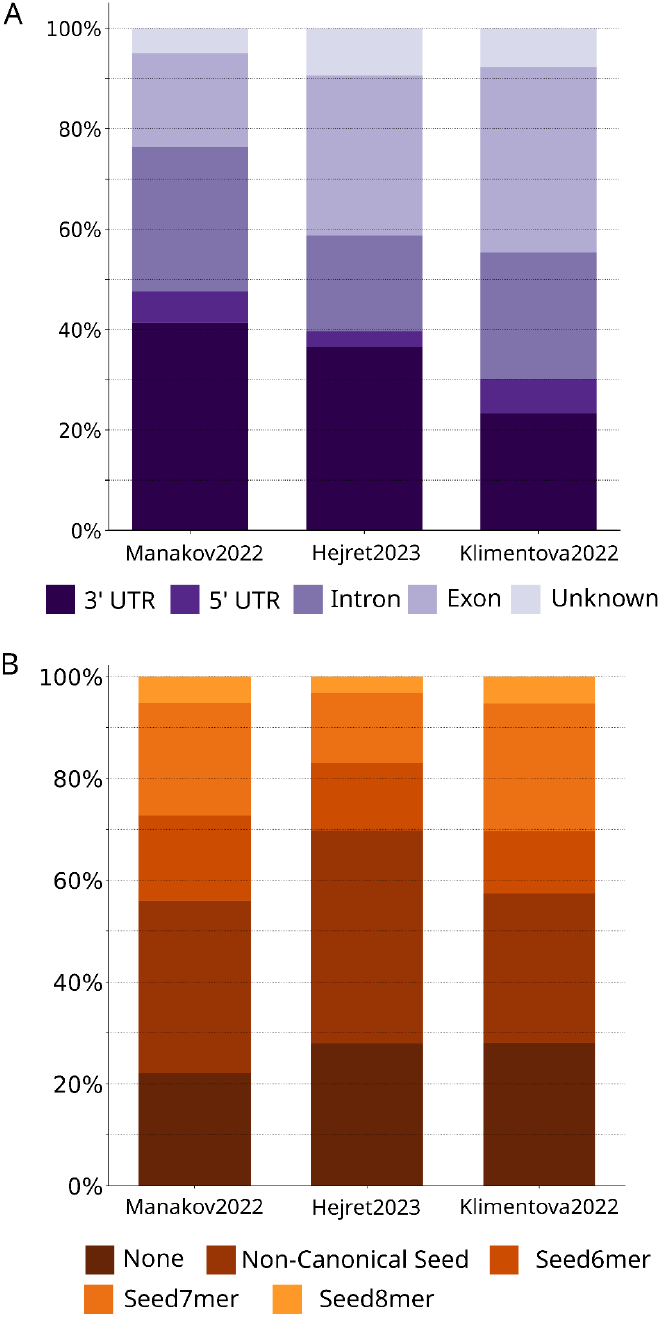
A) Distribution of binding site percentage overlap to genomic element annotations. B) Distribution of miRNA seed types.

Interestingly, most miRNA target prediction methods only take into consideration targets within the 3’UTR, rarely exons, and never introns, despite the binding site distribution seen here. This is a potential blind spot that ignores a large number, potentially a majority, of binding sites from consideration of their effect on the overall regulation of the target messenger RNA.

Given the predominance of seed-like measures for miRNA binding site prioritisation, we explored the prevalence of canonical seed in binding site sequences in the produced datasets (Fig.2B). Canonical seed interactions comprised a minority of each dataset: Manakov2022 (44%), Hejret2023 (30%), and Klimentova2022 (43%). For all three datasets, the Seed7mer was the most prevalent type of canonical seed interaction, at 22%, 14%, and 25% of interactions in Manakov2022, Hejret2023, and Klimentova2022, respectively. Interestingly, interactions lacking exact match for any of the defined seeds comprised only approximately 22% of Manakov2022, and 28% for Hejret2023 and for Klimentova2022. It also appears that the chimeric eCLIP methodology used in Manakov2022 and Klimentova2022 is more effective in capturing canonical seed-type interactions compared to the CLASH method used in Hejret2023, at least for our limited dataset size.

### 3.5 Benchmarking and retraining models

#### 3.5.1 Benchmarking of state-of-the-art miRNA binding site prediction tools on novel datasets

We evaluated multiple state-of-the-art binding site prediction methods including CNNs (TargetScanCnn_McGeary2019, CnnMirTarget_Zheng2022, miRNA_CNN_Hejret2023), ResNets (TargetNet_Min2021, miRBind_Klimentova2022), attention-based models (InteractionAwareModel_Yang2024), and simpler co-folding and seed-based strategies (RNACofold, Seed6mer, Seed7mer, Seed8mer, SeedNonCanonical) on all datasets (Fig. 3, Table 2).

**Table 2.**
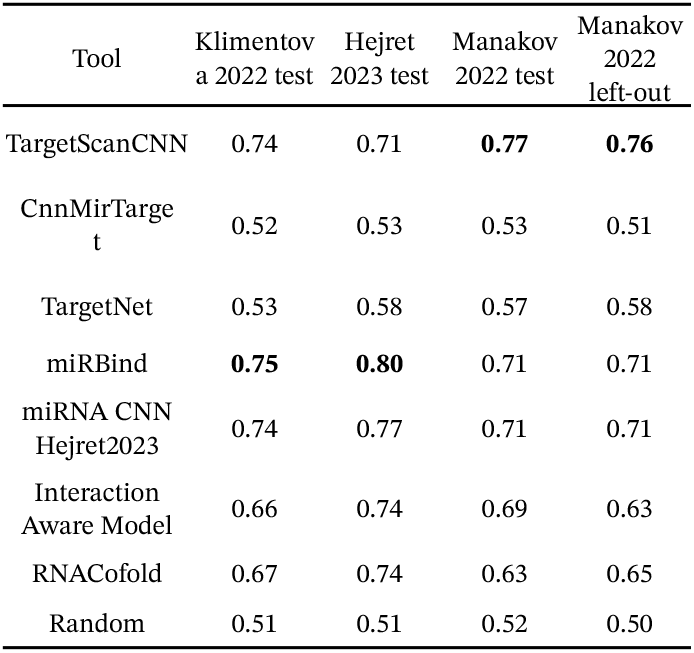
Average Precision Score (APS) from evaluating seven miRNA binding site prediction tools against the four test datasets in miRBench. The best performing model per dataset is highlighted.

| Tool | Klimentov<br>a 2022 test | Hejret<br>2023 test | Manakov<br>2022 test | Manakov<br>2022<br>left-out |
| --- | --- | --- | --- | --- |
| TargetScanCNN | 0.74 | 0.71 | **0.77** | **0.76** |
| CnnMirTarget | 0.52 | 0.53 | 0.53 | 0.51 |
| TargetNet | 0.53 | 0.58 | 0.57 | 0.58 |
| miRBind | **0.75** | **0.80** | 0.71 | 0.71 |
| miRNA CNN<br>Hejret2023 | 0.74 | 0.77 | 0.71 | 0.71 |
| Interaction<br>Aware Model | 0.66 | 0.74 | 0.69 | 0.63 |
| RNACofold | 0.67 | 0.74 | 0.63 | 0.65 |
| Random | 0.51 | 0.51 | 0.52 | 0.50 |

**Figure 3.**
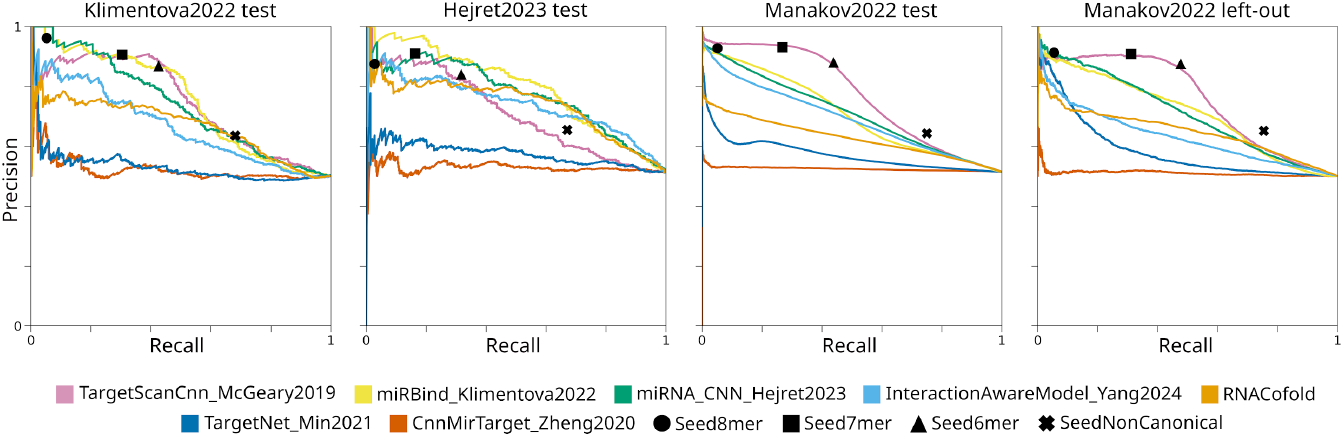
Precision-recall curves. All miRNA binding site prediction tools available on the miRBench package, evaluated on the Klimentova2022 test set, Hejret2023 test set, Manakov2022 test set, and Manakov2022 left-out set.

Within individual test sets, miRBind achieved the highest APS on the Hejret2023 and Klimentova2022 test sets (0.80; 0.75), while TargetScanCnn led on both Manakov test and left-out datasets (0.77, 0.76). When examining cross-dataset performance, TargetScanCnn demonstrated consistent performance across all datasets (0.71-0.77), miRBind showed strong but variable performance (0.71-0.80) with better results on Klimentova2022 and Hejret2023 tests, and miRNA_CNN_Hejret2023 maintained stable performance (0.71-0.77). In contrast, CnnMirTarget and TargetNet showed consistent but lower performance (0.51-0.58) across all datasets, while the InteractionAwareModel displayed moderate performance (0.63-0.74).

Overall, there is not a single state-of-the-art method that consistently outperforms all others in every dataset, leaving the field open for new methods to be developed.

#### 3.5.2 Retrained miRNA_CNN_Hejret2023 benchmarked

To further confirm the importance of unbiased datasets, we retrained the miRNA_CNN_Hejret2023 model proposed by Hejret et al. (2023) exclusively on the new miRNA frequency class unbiased Hejret2023 train set. The intent of this exercise is to understand to which extent the *microRNA frequency class bias* in the original dataset affected the potential of the trained model to learn. We elected to use the miRNA_CNN_Hejret2023 model instead of the better performing miRBind_Klimentova2022 architecture as it is a simpler model consisting of a single convolutional neural network. The miRBind model utilises smaller pilot models and soft labelling of training data, and would need more optimization to be retrained effectively.

We retained unchanged architecture and hyperparameters from the original publication to isolate just the effect of training data quality. We cannot guarantee that this is the optimal model or optimal performance that this model could potentially learn from this dataset. That said, the retrained CNN achieved an APS of 0.86 on the Hejret2023 test set (vs original miRNA_CNN_Hejret2023 model APS=0.77, Table 3) showing significant predictive performance improvement (Fig. 4). This retrained model outperformed the best performing state-of-the-art tools in all test datasets.

**Table 3.** Average Precision Score (APS) of original and retrained models. The best in each dataset is highlighted.

| Tool | Klimentov<br>a 2022 test | Hejret<br>2023 test | Manakov<br>2022 test | Manakov<br>2022<br>left-out |
| --- | --- | --- | --- | --- |
| *Original*<br>miRNA<br>CNN_Hejret2023 | 0.74 | 0.77 | 0.71 | 0.71 |
| *Retrained*<br>on Hejret<br>2023 train set | 0.77 | **0.86** | 0.79 | 0.78 |
| *Retrained*<br>on Manakov<br>2022 train set | **0.84** | 0.84 | **0.84** | **0.81** |

**Figure 4.**
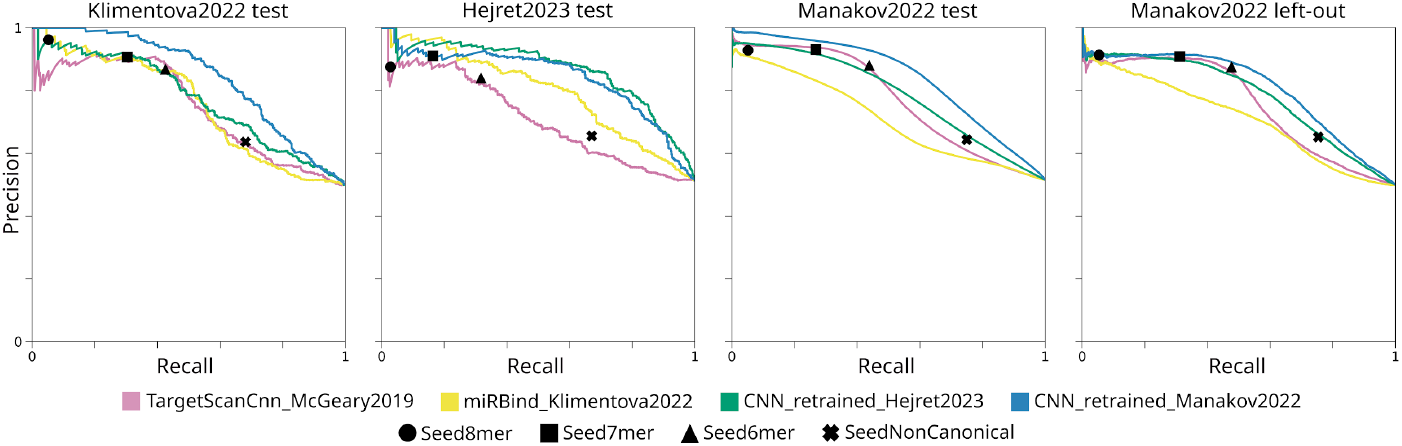
Precision-recall curves. Retrained models (CNN retrained Hejret2023 and CNN retrained Manakov2022), as well as models that previously performed best on any benchmarked dataset (miRBind, TargetScanCnn), evaluated on the Klimentova2022 test set, Hejret2023 test set, Manakov2022 test set, and Manakov2022 left-out set.

We noted that the retrained model performed much better on the Hejret2023 test set (APS = 0.86) compared to the other testing datasets (APS = 0.77-0.79). This leads us to believe that there could be experiment-or dataset-specific elements learnt by the model that still harm full generalisability. The Hejret2023 dataset is the only one produced using the CLASH experimental technique, while all other datasets are produced using the chimeric eCLIP technique. Moreover, the Hejret2023 dataset appears to have retrieved a higher percentage of No Seed (‘None’) and ‘Non-Canonical Seed’ interactions (Fig. 2B). Thus, models trained on this dataset may overestimate the importance of these types of interactions, harming generalizability to other datasets.

Given that some experiment-or dataset-specific elements remain elusive, we advocate for reporting results on all datasets independently in future benchmarking efforts. We expect that incorporating additional experimental datasets into future iterations of miRBench will help uncover experiment– and dataset-specific biases underlying these performance discrepancies.

In addition to the retraining on the Hejret2023 dataset, we also retrained the miRNA_CNN_Hejret2023 model on the larger, unbiased Manakov2022 train set, achieving strong performance across all four test datasets (APS = 0.81 – 0.84, Fig. 4, Table 3). This demonstrates that even with a simple CNN architecture, a well-curated, large training set can yield state-of-the-art results.

#### 3.5.3 Increasing dataset size improves CNN performance with diminishing results

The Manakov2022 train dataset, with 1,253,320 positive examples, is approximately 325 times larger than the Hejret2023 train set (4,084 positive examples). To investigate the influence of training dataset size on model performance, we conducted a series of experiments by retraining the miRNA_CNN_Hejret2023 model on subsets of the Manakov2022 train set, ranging from 100 to 1,253,320 positive examples.

As illustrated in Figure 5, model performance, measured by APS, increased logarithmically with the number of training examples.

**Figure 5.**
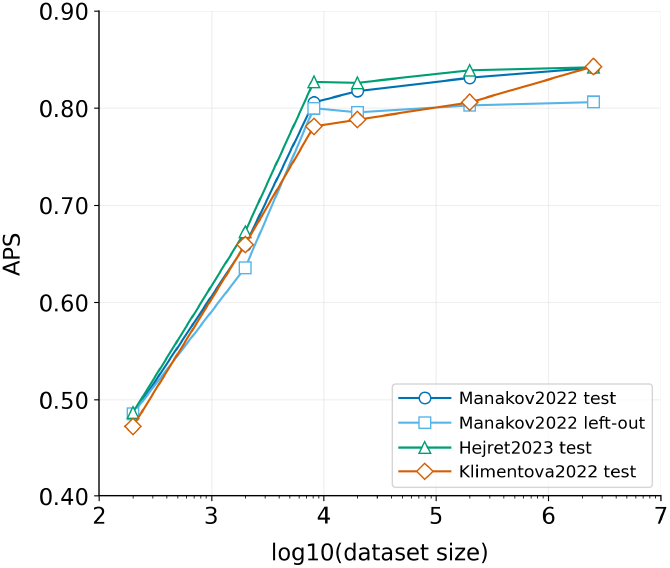
Average precision score against training dataset size for retrained model.

However, this improvement plateaued once the training set size reached approximately 4,084 positive examples, the size of the corrected Hejret2023 train set. Further increasing the dataset size beyond this point yielded only marginal gains, with performance stabilizing at an APS of approximately 0.84 for the three test sets and 0.81 for the Manakov2022 left-out dataset. These findings suggest that while larger datasets are generally beneficial, there are diminishing returns beyond a certain threshold, at least for this specific model architecture.

While our results demonstrate the effectiveness of a relatively simple CNN architecture, it is conceivable that larger, more complex models could extract further insights and achieve even higher performance from these extensive datasets. However, exploring the interplay between model complexity and very large dataset sizes is beyond the scope of this current work, and we encourage further investigation in this area by other research groups.

## 4. Conclusion

This study introduces miRBench, a novel benchmark framework for evaluating miRNA binding site prediction methods, and identifies a critical and previously unreported bias in existing datasets: miRNA frequency class bias. We demonstrate that this bias, stemming from the conventional approach to generating negative training examples, artificially inflates the performance of several state-of-the-art prediction models. To address this issue, we developed a new methodology for negative example generation that effectively mitigates this bias, ensuring a more balanced representation of miRNA families in both positive and negative classes.

Using this improved methodology we curated several new datasets, including the Manakov2022 dataset which is derived from a novel experimental technique and is orders of magnitude larger than any previously available miRNA binding site interaction dataset. Our comprehensive benchmarking on these datasets revealed that while several current methods perform reasonably well, no single method consistently outperforms the others across all datasets. Importantly, we show that a simple convolutional neural network model, when retrained on our unbiased datasets, surpasses the performance of all existing methods on these benchmarks, highlighting the profound impact of dataset quality on model performance.

Furthermore, our investigation into the relationship between dataset size and model performance revealed that while increasing the number of training examples generally improves performance, this improvement plateaus beyond a certain threshold, at least for the CNN architecture employed in this study. This suggests that dataset quality, particularly the absence of biases, is as crucial as, if not more important than, dataset size. The consistently lower performance of models on the Manakov2022 ‘left-out’ dataset, which contains unique miRNA families, underscores the importance of diverse and representative training data for achieving true model generalizability.

To facilitate future research, we provide miRBench, an open-source Python package that offers easy access to the curated datasets, encoders, and implementations of state-of-the-art prediction models. We believe that miRBench, with its unbiased datasets and user-friendly interface, will serve as a valuable resource for the development and evaluation of more accurate and robust miRNA binding site prediction methods. Future work should focus on exploring more complex model architectures capable of leveraging the full potential of large-scale datasets, as well as further investigation into the dataset– and experiment-specific factors that contribute to the remaining performance discrepancies. Ultimately, a deeper understanding of miRNA-target interactions, driven by rigorous benchmarking and unbiased datasets, will be crucial for unlocking the full potential of miRNAs in both basic biology and clinical applications.

## Data availability

The latest version of the miRBench package is freely available on GitHub at https://github.com/katarinagresova/miRBench/tree/v1.0.0, and has a permanent version deposited on Zenodo at https://zenodo.org/records/14652776. The Python package is also distributed through the Python Package Index (PyPI) at https://pypi.org/project/miRBench/1.0.0/.

The datasets produced were deposited on Zenodo at https://zenodo.org/records/14501607, so they are easily accessible by the miRBench Python package. Code for processing datasets is available at https://github.com/BioGeMT/miRBench_paper/tree/v1.0.0

## Acknowledgements

The authors would like to thank E. Maragkakis, M. Ciach, A. Balestrucci, and V. Martínek for their constructive criticism of the manuscript.

## Footnotes

1 https://github.com/ML-Bioinfo-CEITEC/HybriDetector/blob/main/ML/Datasets/miRNA_train_set.tsv

2 https://github.com/ML-Bioinfo-CEITEC/HybriDetector/blob/main/ML/Datasets/miRNA_test_set_1.tsv

3 https://bitbucket.org/bipous/miraw_data/src/master/ (accessed on 16-Dec-2024)

4 miraw_data/PLOSComb/Data/ValidTargetSites/allTrainingSites.txt

5 miraw_data/PLOSComb/Data/TestData/balanced10/randomLeveragedTestSplit_0.csv

6 https://github.com/enasequence/enaBrowserTools (accessed May 2024)

7 https://github.com/ML-Bioinfo-CEITEC/HybriDetector

8 https://github.com/ML-Bioinfo-CEITEC/HybriDetector/tree/fix_clustering

9 https://zenodo.org/records/14501607/files/AGO2_eCLIP_Manakov2022_full_dataset.tsv.gz

10 https://github.com/ML-Bioinfo-CEITEC/miRBind/blob/main/Datasets/AGO2_eCLIP_Klimentova22_full_dataset.tsv

11 https://github.com/ML-Bioinfo-CEITEC/HybriDetector/blob/main/ML/Datasets/AGO2_CLASH_Hejret2023_full_dataset.tsv

12 https://github.com/BioGeMT/miRBench_paper/tree/v0.2.0/code/post_process

